# Peripheral nerve injury results in a biased loss of sensory neuron sub-populations

**DOI:** 10.1101/2023.11.14.566863

**Authors:** Andrew H. Cooper, Allison M. Barry, Paschalina Chrysostomidou, Romane Lolignier, Heather F. Titterton, David L. Bennett, Greg A. Weir

## Abstract

There is a rich literature describing loss of dorsal root ganglion (DRG) neurons following peripheral axotomy, but the vulnerability of discrete subpopulations has not yet been characterised. Furthermore, the extent and even presence of neuron loss following injury has recently been challenged. In this study, we have used a range of transgenic recombinase driver mouse lines to genetically label molecularly defined subpopulations of DRG neurons and track their survival following traumatic nerve injury. We find that spared nerve injury (SNI) leads to a marked loss of cell containing DRG-volume and a concomitant loss of small diameter DRG neurons. Neuron loss occurs unequally across subpopulations and is particularly prevalent in non-peptidergic nociceptors, marked by expression of Mrgprd. We show that this subpopulation is almost entirely lost following SNI and severely depleted (by roughly 50%) following sciatic nerve crush. Finally, we used an *in vitro* model of DRG neuron survival to demonstrate that non-peptidergic nociceptor loss is likely dependent on the absence of neurotrophic support. Together, these results profile the extent to which DRG neuron subpopulations can survive axotomy, with implications for our understanding of nerve injury-induced plasticity and pain.

## Introduction

Dorsal root ganglion (DRG) neurons represent a molecularly and functionally heterogeneous population. Under normal conditions, this diversity contributes to the ability of the somatosensory nervous system to detect a myriad of sensory stimuli that result in the perceptions of touch, temperature, itch, and pain. Following nerve injury, physiological changes in DRG neurons lead to hyperexcitability [56], which is a key pathological driver of neuropathic pain [20,62]. Concomitant molecular changes in discrete sub-populations also occur, and these have recently been comprehensively described in single-cell [36,43] and sub-population-specific sequencing studies [3]. These studies describe a transient and generalised reduction in the expression of subpopulation specific genes following nerve injury [3,36,43].

In addition to molecular changes, there is a rich literature describing frank loss of DRG neurons following traumatic nerve injury in experimental rodent models [24,49,52,55] and human patients [47,65]. In rodents, most studies support a preferential loss of small cells that give rise to unmyelinated fibres [52] but some contrasting studies describe preferential loss of large cells [6] or loss of cells of all sizes [45]. Variation is evident across studies in terms of experimental species, age, type of injury, and quantification methods. Vestergaard *et al* reported a loss of 22% of DRG neurons two weeks after transection of the spinal nerve 7mm distal to L5 DRG in rat, which rose to a 35% loss by 7 weeks [55]. In contrast, Tandrup *et al* described a delayed loss (no significant loss before 8 weeks after transection) of 43% of small DRG neurons by 32 weeks following “mid-thigh” sciatic nerve lesion in rat, which suggests that the site of nerve lesion is important [52]. Shi *et al* used stereological counting methods to identify a 54% loss of DRG neuron number 4 weeks after “mid-thigh” sciatic nerve transection in C57BL/6 mice [49]. Estimates for the degree of loss following commonly used nerve injury paradigms (e.g. spared nerve injury (SNI) and sciatic nerve crush (SNC)) are not available and [46][43] because of the neurochemical changes following injury and the loss of subpopulation marker gene expression [5,43,49], the vulnerability of molecularly defined subpopulations has not been characterised. Moreover, more recent studies have even cast doubt on the extent, or presence of DRG neuron death following nerve injury. One study which developed a deep learning approach to assess rat DRG cellular plasticity found no loss of neurons up to two weeks post-SNI [48], while another observed no loss of genetically labelled damaged DRG neurons two months after SNC [43].

The issue of whether neuron loss occurs, and if so, in what subpopulations, is important. It will likely have implications for our understanding of reinnervation and functional recovery in patients. Furthermore, better insight will provide critical context for those investigating the plasticity that occurs following nerve injury and may inform therapeutic targeting of sensory neuron populations.

An expanding repertoire of transgenic recombinase driver lines now makes it possible to permanently label DRG neuron subpopulations and study their fate in rodent nerve injury paradigms. The aim of this study was to use this technology to characterise neuron loss after nerve injury and to test the hypothesis that loss is not equally distributed across the molecular populations.

## Methods

### Animals

Mice were group-housed in humidity and temperature-controlled rooms with free access to food and water, on a 12-hour light-dark cycle, and with environmental enrichment. Animal procedures were performed under a UK Home Office Project Licence and in accordance with the UK Home Office (Scientific Procedures) Act (1986). All studies were approved by the Ethical Review Process Applications Panel of the University of Glasgow or Oxford and conform to the ARRIVE guidelines. Experiments were performed on a mix of adult male and female mice, aged 7-16 weeks at the beginning of experiments. Details of transgenic lines are provided in **Table 1** [1,4,13,16,32,38,39,50,58,71]. Tamoxifen was administered by i.p. injection of 20 mg/mL tamoxifen (Sigma Aldrich) dissolved in wheat germ oil (doses described in **Table 1**). There were two instances where animals were excluded from data analysis: One Thy1-CFP died of unknown causes not related to the procedure and prior to the experimental endpoint, and one MrgD^CreERT2^;Ai32 exhibited no fluorophore expression and was therefore deemed to have been incorrectly genotyped. Group sizes were based on the extent of neuronal loss 28d following sciatic nerve transection identified by Shi *et al*. [49]: given α = 0.05, power = 0.8 and an effect size of 4.81, power analysis projects that a group size of 3 mice would be needed.

**Table 1.** Transgenic lines used in the study.

| Used name | Full name | Ref | Source | Tamoxifen regime |
| --- | --- | --- | --- | --- |
| Atf3<sup>CreERT2</sup> | Atf3<sup>tm1.1(cre/ERT2)Msra</sup> | [8] | Gift: Dr Franziska Denk | 50 mg/kg on days 0, 3 and 7 after surgery. |
| Avil<sup>FlpO</sup> | Avil<sup>tm1(flpo)Ddg</sup> | [55] | Gift: Prof David Ginty | N.A. |
| MrgD<sup>CreERT2</sup> | Mrgprd<sup>tm1.1(cre/ERT2)Wql</sup> | [25] | The Jackson Laboratory (#031286) | General: 1x 50 mg/kg in adulthood, (> 1 week prior to experiment).<br>3D volumetric analysis: 5x i.p. (0.5 mg/animal/day), beginning between P10-P17 |
| MrgD<sup>Chr2-YFP</sup> | Mrgprd<sup>tm4.1(COP4)Mjz</sup> | [43] | Mutant Mouse Resource & Research Centers (036112-UNC) | N.A. |
| Calca<sup>CreERT2</sup> | Calca<sup>tm1.1(cre/ERT2)Ptch</sup> | [37] | Gift: Prof Pao-Tien Chuang | 1x 75 mg/kg in adulthood (> 1 week prior to experiment). |
| Trpm8<sup>FlpO</sup> |  | [4] | Gift: Dr Mark Hoon | N.A. |
| RC::FLTG | B6.Cg-*Gt(ROSA)26Sor<sup>tm1.3(CAG-tdTomato,-EGFP)Pjen</sup>* | [26] | The Jackson Laboratory (#026932) | N.A. |
| Ai32 | B6.Cg-*Gt(ROSA)26Sor<sup>tm32(CAG-COP4\*H134R/EYFP)Hze</sup>* | [21] | The Jackson Laboratory (#007909) | N.A. |
| Thy1-CFP | B6.Cg-Tg(Thy1-CFP)23Jrs/J | [11] | The Jackson Laboratory (#003710) | N.A. |
| Th<sup>CreERT2</sup> | Th<sup>tm1.1(cre/ERT2)Ddg/J</sup> | [1] | Gift: Prof David Ginty ; CAT#:025614 | 1x 50 mg/kg in adulthood (> 2 week prior to experiment) |

### Spared nerve transection and crush surgeries

Spared nerve injury (transection of the common peroneal and tibial branches of the sciatic nerve; SNI_trans_) and common peroneal and tibial crush injury (SNI_crush_), in which nerve axons were severed but the epineurium remained intact, were performed as previously described [12]. Anaesthesia was induced with 3-5% isoflurane and then maintained at 1.5-2% as required. Analgesia, consisting of carprofen (10 mg/kg) and buprenorphine (0.05 mg/kg), (Glasgow) or carprofen (5 mg/kg) and local bupivacaine (2 mg/kg) (Oxford) was provided perioperatively. The left hindpaw was secured with tape in hip abduction and the operative field (lateral surface of the thigh) was shaved. Ophthalmic ointment was applied to the eyes and the shaved area was swabbed with chlorhexidine solution. A longitudinal incision was made in the skin at the lateral mid-thigh. Using blunt dissection an opening was made through the biceps femoris, exposing the sciatic nerve and the three peripheral branches (sural, tibial, and common peroneal nerves). For SNI_trans_, the common peroneal and tibial nerves were ligated using 6-0 Vicryl suture (Ethicon, USA) and a 1-2 mm piece distal to the suture was removed using spring scissors. For SNI_crush_, the exposed tibial and common peroneal nerves were clamped using a pair of fine haemostats (Fine Science Tools, Germany) closed to their second clip, leaving the nerve branches intact but translucent. The muscle was closed with one 6-0 Vicryl suture (Ethicon, USA), and the skin incision closed with one 10 mm wound clip (Alzet, USA). Animals were monitored daily for self-mutilation, and no animals required sacrifice due to tissue damage.

### FastBlue tracer injections

Mice were briefly anaesthetised during the procedure, induced with 3-5% isoflurane then maintained at 1.5-2% as required. MrgD^CreERT2^;Ai32 mice were used for data presented in Figure 2, without revealing MrgD-YFP signal. Hindlimbs were taped with the plantar surface of the paw facing up, and a custom, 26G removable needle with 30° bevel, attached to a 25 µL Hamilton syringe, was inserted between the two distal-most footpads, towards the medial aspect of the hindpaw. The needle was then rotated 90°, so the bevel faced medially. 4 µL FastBlue (FB; 2% in sterile PBS; CAS# 73819-41-7; Polysciences Inc., Warrington, PA USA) per paw was then slowly injected, and the needle left in place for 10s, before rotating and carefully retracting to avoid backflow of FB along the needle track. This prevented the FB bolus from contacting the sural innervation territory of the lateral hindpaw, restricting it largely to the tibial innervation territory of the glabrous hindpaw skin.

### Immunohistochemistry and image acquisition

Mice were anaesthetised with an overdose of pentobarbital (20 mg) and transcardially perfused with fixative containing 4% formaldehyde. L3 to L5 DRG were removed and post-fixed for a further 2 hours, cryoprotected in 30% sucrose overnight, and then embedded in optimal cutting temperature media (OCT; Tissue Tek, USA). DRG were sectioned on a Leica CM1950 cryostat at 30 µm, with every section collected serially on 5 Superfrost Plus slides (VWR, Lutterworth, UK) and each slide containing 1 in every 5 sections (4-7 sections per slide). One slide per DRG was selected at random and was washed with PBS, before being incubated with appropriate primary antibodies (**Table 2**) diluted in 5% normal donkey serum and 0.3% Triton X-100 in PBS for 3 days at 4°C. After PBS washes, slides were incubated with appropriate secondary antibodies (**Table 2**) in the same PBS/NDS/Triton-X100 solution as for primaries, overnight at room temperature. Slides were washed and coverslipped with VectaShield Vibrance Hardset mounting media (Vector Labs, CA, USA), with DAPI included in mounting media where FastBlue-labelled cells were not being examined. Sections were imaged using a Zeiss LSM900 Airyscan confocal microscope equipped with 405, 488, 561 and 640 nm diode lasers. Full thickness, tiled, confocal image stacks with a 2-3 µm interval in the Z-axis were obtained through a 20× dry lens (0.8 NA) with the confocal aperture set to 1 Airy unit or less. All image capture was performed using Zen Blue Edition software (Carl Zeiss Microscopy GmbH, Germany), and analyses were performed using Zen Blue or FIJI [44].

**Table 2.** Primary and secondary antibodies used in the study.

| **Antibody** | **Source** | **Identifiers** | **Working dilution** |
| --- | --- | --- | --- |
| Anti-GFP (Chicken polyclonal) | Abcam plc., Cambridge, UK | Cat#: ab13970<br>RRID: AB_300798 | 1:1000 |
| Anti-NeuN (Guinea pig polyclonal) | Synaptic Systems, Göttingen, Germany | Cat#: 266004<br>RRID: AB_2619988 | 1:500 |
| Anti-mCherry (Rat monoclonal) | Invitrogen; Thermo Fisher Scientific, UK | Cat#: M11217 RRID: AB_2536611 | 1:500 |
| Anti-Atf3 (Rabbit polyclonal) | Novus Biologicals, Minneapolis, MN, USA | Cat#: NBP1-85816<br>RRID: AB_11014863 | 1:500 |
| Anti-NF200 (Rabbit polyclonal) | Sigma-Aldrich, Saint Louis, MO, USA | Cat#: N4142<br>RRID: AB_477272 | 1:1000 |
| Anti-TrkA (Rat polyclonal) | R&D Systems, Minneapolis, MN, USA | Cat#: AF1056<br>RRID: AB_2283049 | 1:500 |
| anti-TDP43 (Rabbit polyclonal) | Abcam plc., Cambridge, UK | Cat#: ab133547<br>RRID: AB_2920621 | 1:100 |
| Anti-RFP (Mouse monoclonal) | Thermo Fisher Scientific, UK | Cat#: MA5-15257<br>RRID: AB_10999796 | 1:200 |
| Anti-RFP (Chicken polyclonal) | Sigma-Aldrich, UK | Cat#: AB3528<br>RRID: AB_11212735 | 1:200 |
| Alexa Fluor 488 Donkey Anti-Chicken IgY (Donkey polyclonal) | Jackson ImmunoResearch, PA, USA | Cat#: 703-545-155<br>RRID: AB_2340375 | 1:500 |
| Alexa Fluor 647 Donkey Anti-Guinea pig IgG (Donkey polyclonal) | Jackson ImmunoResearch, PA, USA | Cat#: 706-605-148<br>RRID: AB_2340476 | 1:250 |
| Rhodamine Red-X Donkey Anti-Rat IgG (Donkey polyclonal) | Jackson ImmunoResearch, PA, USA | Cat#: 712-295-153<br>RRID: AB_2340676 | 1:100 |
| Alexa Fluor 647 Donkey Anti-Rabbit IgG (Donkey polyclonal) | Jackson ImmunoResearch, PA, USA | Cat#: 711-605-152<br>RRID: AB_2492288 | 1:250 |
| Rhodamine Red-X Donkey Anti-Rabbit IgG (Donkey polyclonal) | Jackson ImmunoResearch, PA, USA | Cat#: 711-295-152<br>RRID: AB_2340613 | 1:100 |
| Alexa Fluor 546 Goat Anti-Chicken IgG (Goat polyclonal) | Thermo Fisher Scientific, UK | Cat#: A11040<br>RRID: AB_2534097 | 1:400 |
| Alexa Fluor 488 Goat Anti-Rabbit IgG (Goat polyclonal) | Thermo Fisher Scientific, UK | Cat#: A11008<br>RRID: AB_143165 | 1:400 |
| Alexa Fluor 546 Donkey Anti-Mouse IgG (Donkey polyclonal) | Thermo Fisher Scientific, UK | Cat#: A10036<br>RRID: AB_2534012 | 1:400 |

### Image analysis

During all image quantification, the experimenter was blind to experimental groups. For quantification of total numbers of cells within the DRG, a modified optical dissector stereological method was used [11,18,46] (applies to data presented in Fig 1 A-F and Fig 3 A and B). In case of tissue shrinkage during processing, the mean thickness (*t*) of each section on one slide (i.e., 1 in 5 sections) was calculated by taking the mean of the thickest and thinnest cell-containing regions (i.e., not fibre tract-containing regions) of the section (NB: no optical correction to thickness was applied; given the use of a dry lens, this value will not reflect actual section thickness, though this was kept consistent throughout the study). The cell-containing, cross-sectional area (*a*) was then calculated, using the middle optical section from the series and drawing around the cell-containing area. Section volume (*V_sec_*) was then calculated:

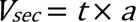

Using the Cavalieri principle, the cell-containing volume of the DRG was calculated [11]:

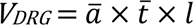

Where 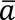 = mean cell-containing cross-sectional area, 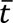 = mean section thickness, and *l* = “length” of the DRG (determined from the total number of sections collected). The number of neurons per section (*N_sec_*) was quantified, including only neurons with a visible nucleus (in the NeuN channel), excluding cells with a nucleus visible within the top frame of the Z-stack, and including any neurons with a nucleus visible in any other field within Z-stack, including the bottom frame of Z-stack. All immunostained sections were counted when examining MrgD-YFP+ neuronal profiles, and for Avil^FlpO^;Atf3^CreERT2^;RC::FLTG, 2 sections per DRG were randomly chosen (via random number generator within Microsoft Excel) and quantified. The cell density, or number of cells per unit vol (*N*_v_) was then calculated:

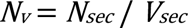

Finally, the total number of cells per DRG (*N_DRG_*) was calculated:

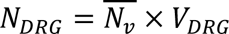

For quantification of the proportion of FB-labelled cells co-labelled with afferent subpopulation markers, initially, the total number of FB-filled neuronal cell profiles with a visible nucleus anywhere within the section was counted, with the observer blind to other channels. The other channel was then revealed, and instances of co-labelling were quantified. No stereological correction was applied, given that the similar size of neuronal nuclei would prevent over-counts of large neurons, and that no comparisons of total number of labelled cells were made. For soma area analyses, the area of neuronal soma expressing the appropriate marker was measured in the optical section within the Z-stack in which that neuron was at its largest, by drawing around the perimeter of the neuron in Fiji/ImageJ v2.14.0/1.54f.

### Tissue clearing and 3D volumetric analyses

DRG were extracted from animals 4 weeks post-SNI_trans_ for whole DRG analyses. Here, tissue was extracted from a combination of MrgD^CreERT2^;Ai14, Th^CreERT2^;Ai14, and Calca^CreERT2^;Ai14 lines (mixed sex)[3]. One month after SNI_trans_, animals were transcardially perfused with sterile saline followed by fixative containing 4% formaldehyde. Ipsilateral and contralateral L4 DRG were removed and post-fixed for 24 hours on a shaker at room temperature before being washed in PBS and stored at -80 in CI-VM1 (35% DMSO, 35% Ethylene Glycol in PBS) until clearing. Tissue clearing was then performed as previously described [66]. Briefly, tissue was exposed to a gradient of 1-propanol containing 0.3% tirethylamine (30, 50, 75, 90, 95, 100, 100%), and washed in this solution at 37°C for 24 hours. Tissue was then rehydrated in PBS and labelled with primary antibodies for 1 week at 37°C (mouse anti-TDP43 and 2x anti-RFP, **Table 2**). Tissue was washed for 24 hours and incubated with appropriate secondary antibodies (**Table 2**) for another week at 37°C. Tissue was subsequently washed for 24 hours, dehydrated again in increasing concentrations of 1-propanol containing 0.3% triethylamine, and mounted in benzyl alcohol with benzyl benzoate (1:2 ratio) containing 0.3% triethylamine on glass slides with silicone spacers. Imaging was performed on an Olympus spinning disk confocal microscope at 20x, with 2 µm z-steps. Tissue was stored at 4°C for ∼ 16 months prior to imaging and so only tissue which remained transparent at this time was used for downstream analyses. Volumetric analyses were performed via Imaris using the “spots” feature with region growing (to allow for different sized spots), background subtraction, and point spread function (PSF) elongation (standard 2*XY). Initial spot diameters were set based on MrgD^CreERT2^;Ai14 nuclear size (as labelled by RFP). Spot classification was then performed blind by adjusting the quality threshold to balance detection in superficial and deep tissue. This step was necessary due to differences in tissue quality after long term storage. Any labelled spots in the adjacent nerve were then deleted (e.g., labelled Schwann cells or debris). Count and volumetric data were then exported for analysis in R. Data were filtered for very small (<5 µm^3^) and very large (>2000 µm^3^) spots to further remove any debris, labelled satellite glia or doublets within the ganglia. In both cases, these filters were approximate and do not exclude the possibility that some spots correspond to either class in the final dataset. The upper limit of "small" DRG nuclei size category was defined as the upper bound of 32 easily identifiable MrgD+ nuclei (258 µm^3^). The boundary between “medium” and “large” bins (400 µm^3^) was less clearly defined in the samples, and was therefore set as the approximate mid-point of the volume distribution. A combined size category for all nuclei greater than 258 µm^3^ was also examined, and results mirrored those of “medium” and “large” bins.

### Gene Ontology

Gene Ontology (GO) term analyses were performed on previously published mouse subtype RNA-seq after SNI (GSE216444 [3]). Here, subtype-specific bulk RNA-seq was performed on 5 transgenic mouse lines through reporter labelling and FACS. STAR was used to map reads to the GRCm38 (mm10) Mouse Genome [14] and Samtools was used to sort, index, and merge Binary Alignment Map (BAM) files in line with published reports [28]. Quality control mirrored Barry et al. (2023). Downstream analyses were performed using DESeq2 on grouped male and female samples [31]. For differentially expressed genes (DEGs; FDR < 0.05, LFC >1), GO analyses were performed using the Wallenius method via goSeq (R) [68]. Here, significantly regulated terms related to cell death and apoptosis with more than 10 genes were examined. The filtered count data of expressed, non-DEG genes were used as a background.

### Dorsal root ganglion culture

DRG were dissected from MrgD^CreERT2^;Ai32 mice >1 week after dosing with tamoxifen, and enzymatically digested at 37°C for 80 min in dispase type II (4.7 mg/ml) plus collagenase type II (4 mg/ml) (Worthington Biochemical), as described previously [62]. Mechanically dissociated cells were plated onto laminin/poly-D-lysine (R&D Systems) treated coverslips in complete Neurobasal® Plus medium [Neurobasal® Plus media supplemented with 2% (v/v) B27 Plus, 1% N2, 1% Glutamax and 1% antibiotic-antimycotic (ThermoFisher Scientific)]. Mouse nerve growth factor (50 ng/ml; NGF, PeproTech) and 10 ng/ml glial-derived neurotrophic factor (GDNF, PeproTech) were added to the media under some conditions. Cytosine β-D-arabinofuranoside (AraC, 4µM) was added to media for 24hrs the day after plating to reduce proliferation of non-neuronal cells. Media was refreshed three times per week thereafter. Cultures were fixed for 10 min at room temperature with 4% PFA and subsequently processed by immunocytochemistry (described above).

### Statistical analysis

Data are expressed as mean ± SEM unless otherwise specified, and *P* values of less than 0.05 were considered significant. Power calculations were performed using G*Power 3.1.9.7 [15]. Quantitative Venn diagram was created using BioVenn [25]. All other statistical analyses were performed in Prism 10 (GraphPad Software Inc., USA) or R using paired t-tests, one, or two-way RM ANOVAs, where appropriate. Normality was assessed by Shapiro–Wilk test. If a main ANOVA effect was significant, then Šídák’s or Tukey’s multiple comparisons tests were performed. To compare population distributions of soma cross-sectional area or volume, Kolmogorov-Smirnov tests were performed.

## Results

### Peripheral nerve injury induces a loss of small neurons from the dorsal root ganglion

To assess the gross loss of neurons from DRG following nerve injury, we generated the Avil^FlpO^;Atf3^CreERT2^;RC::FLTG mouse line in which naïve and axotomized sensory neurons were differentially labelled. In this mouse line, all neurons express tdTomato (Flp-dependent) in the naïve state, and switch to expressing GFP upon axonal damage and concurrent tamoxifen treatment (Flp-and Cre-dependent) (**Fig. 1A-B**). Following pilot experiments to optimise tamoxifen dosing regimen, this approach was both highly efficient and specific (with the caveat that it was necessary to wait for several days after nerve injury for Cre-induced GFP expression): 14 days after SNI_trans_ surgery, GFP was expressed by 99.1 ± 0.6% of Atf3-expressing ipsilateral L4 DRG neurons, while we observed GFP in only 4.6 ± 0.7% of contralateral DRG neurons (**Fig. S1A-D**). We then used a stereological approach to quantify the total number of neurons in L4 DRG ipsilateral to injury 1, 2, 4 and 8 weeks after SNI_trans_. One week after SNI_trans_, we observed 7809 ± 153 neurons per DRG; this approximately halved by 8 weeks post-injury to 3963 ± 410 neurons per DRG (**Fig. 1C**). Separating analysis into intact versus axotomized afferents revealed that only axotomized afferents were lost, with no difference observed in numbers of intact afferents (**Fig. 1D**). Between 1 and 8 weeks after injury, we observed a 61.0 ± 7.0% decrease in the number of GFP+ neurons. This loss of injured afferents resulted in a loss of neuron-containing (i.e., excluding white matter regions) DRG volume (**Fig. 1E**), but not neuron density (**Fig. 1F**). Population distributions of the cross-sectional area of nucleated, tdTomato-expressing cell profiles were not significantly different at 1 versus 8 weeks post-SNI_trans_, in contrast to GFP-expressing/injured afferents, in which a loss of a population of small afferents at 8 weeks post-injury was observed (**Fig. 1G**).

**Figure 1.**
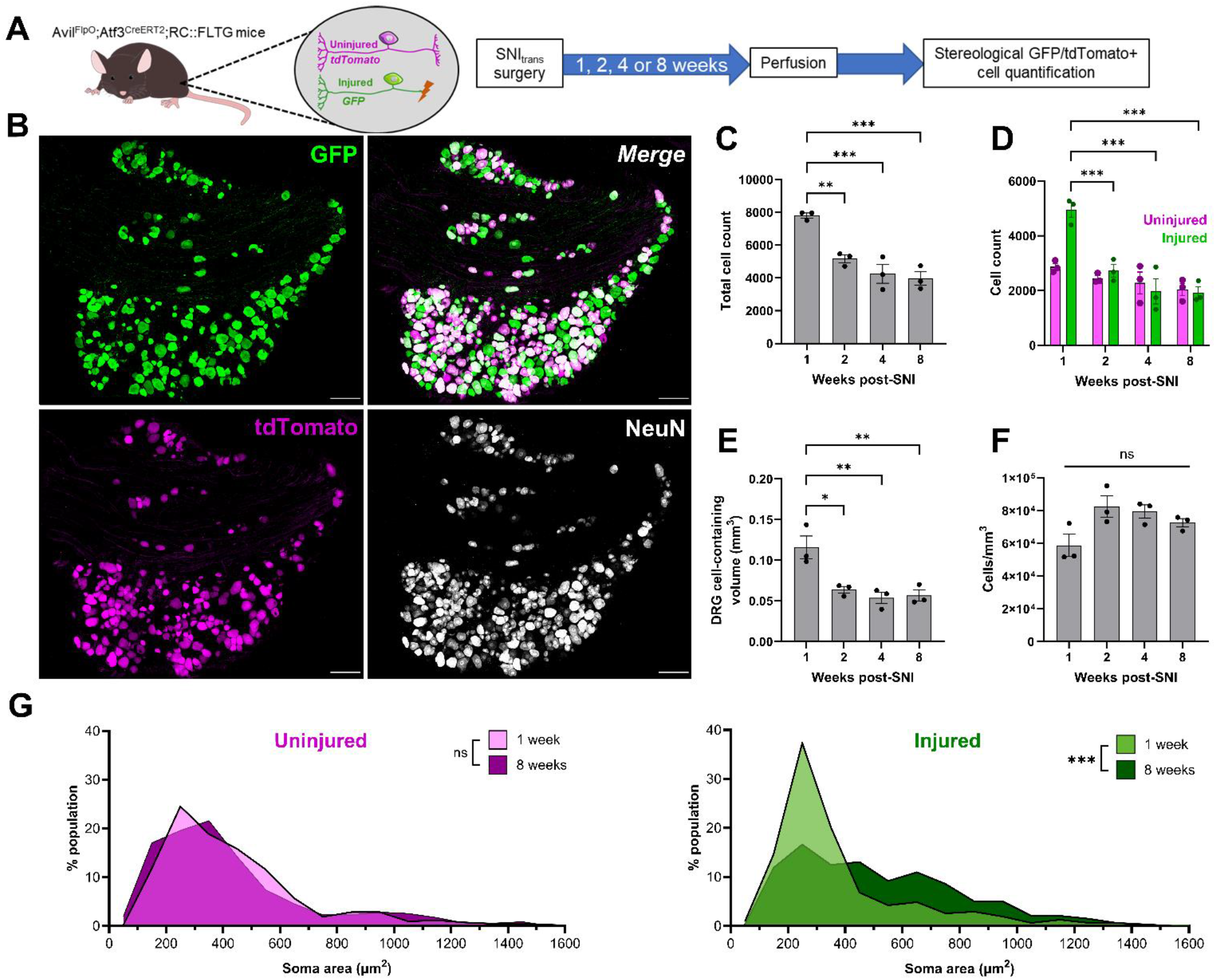
SNI_trans_ induces death of small primary afferent neurons, accompanied by a reduction in volume, not cell density, of the dorsal root ganglion. (A) Approach to differentially label intact afferents with tdTomato and damaged afferents with GFP after peripheral nerve injury using the Avil^FlpO^;Atf3^CreERT2^;RC::FLTG mouse line, and schematic of experimental timeline. (B) Representative image of GFP, tdTomato and NeuN expression in an L4 DRG, 2 weeks after SNI_trans_. Scale bars = 100 µm. (C-D) Stereological quantification of the total number of DRG neurons (C) or number of axotomized and intact neurons (D) in the L4 DRG 1, 2, 4 and 8 weeks after SNI_trans_. C: One-way ANOVA with Tukey’s post-tests; F_3,8_ = 21.30, *P* < 0.001. D: Two-way RM ANOVA with Tukey’s post-tests; Timepoint x Colour interaction F_3,8_ = 7.134, *P* = 0.01, *n* = 3. (E) Volume of DRG containing cells (i.e., excluding white matter tracts) following SNI_trans_. One-way ANOVA with Tukey’s post-tests; F_3,8_ = 10.84, *P* = 0.003, *n* = 3. (F) Neuronal density within the DRG following SNI_trans_. One-way ANOVA; F_3,8_ = 4.03, *P* = 0.051, *n* = 3. (G) Population distribution of uninjured and injured afferents by cross-sectional area, 1- and 8-weeks post-SNI_trans_. Kolmogorov-Smirnov tests of cumulative distributions; Uninjured: D = 0.08, *P* = 0.18; Injured: D = 0.32, *P* < 0.001; *n* = 310 to 427 neurons from 3 mice. * *P* < 0.05, ** *P* < 0.01, *** *P* < 0.001.

SNI_trans_ resulted in a mixed population of axotomised and intact afferents within the L4 DRG. Therefore, we developed an approach to restrict our analysis to axotomised afferents, without relying on transgenic labelling, and used this as a complementary approach to confirm our findings. We injected the neuronal tracer FastBlue (FB) into the glabrous, tibial innervation territory of both hindpaws 1 week prior to common peroneal and tibial transection (SNI_trans_) or crush (SNI_crush_) surgeries (**Fig. 2A-B**). FB-uptake was complete across neurons of all sizes by one week (**Fig. S2**) and so this approach allowed us to profile a sample of the axotomized afferents. Both SNI_trans_ (**Fig. 2C**) and SNI_crush_ (**Fig. 2D**) injuries resulted in a rightward shift in population distributions of the cross-sectional area of nucleated, FB-labelled DRG neurons when compared to contralateral DRG.

**Figure 2.**
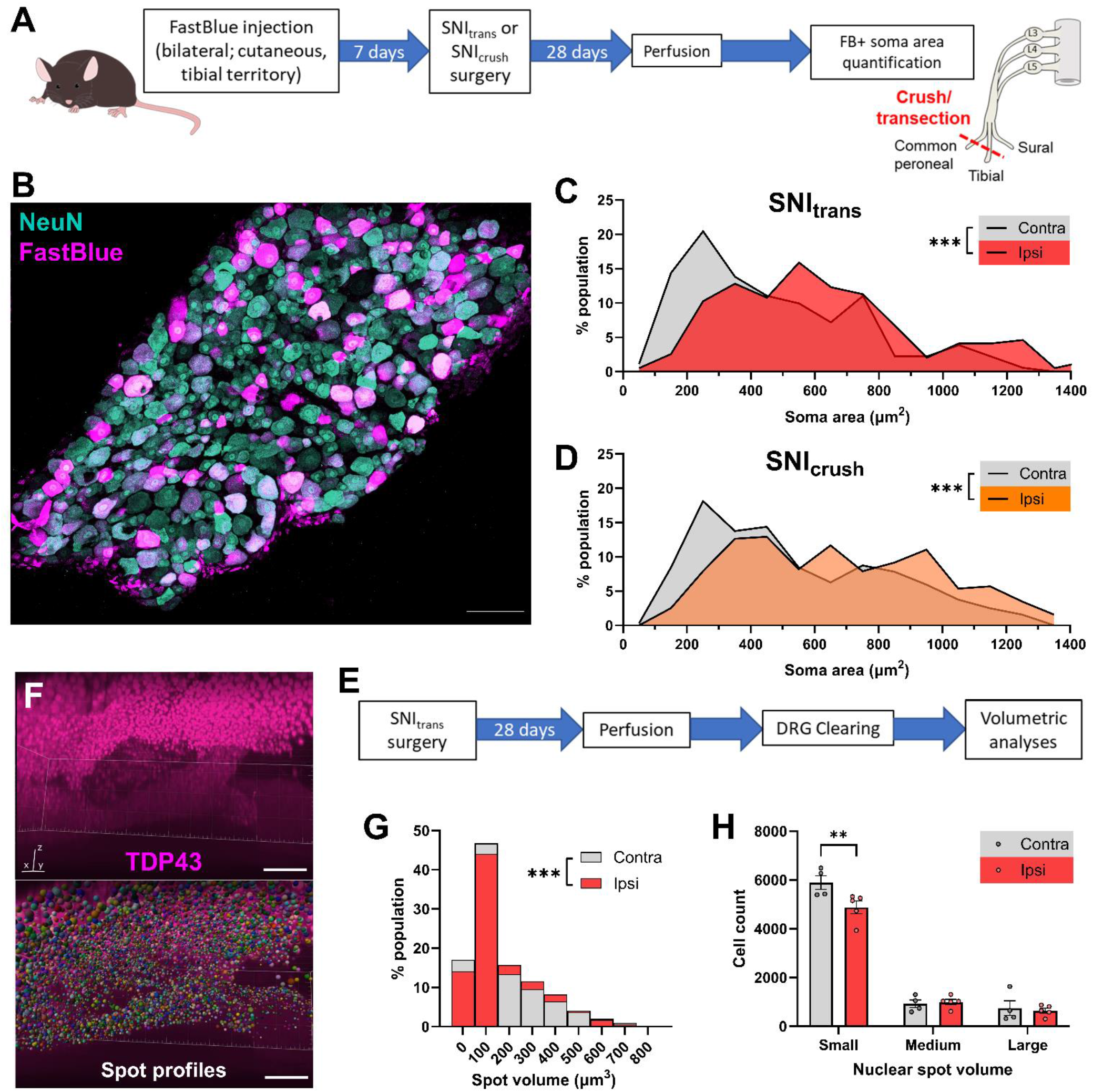
Spared nerve crush and transection lead to a loss of small DRG neurons. (A) Approach to restrict analysis to damaged afferents: a subcutaneous injection of the tracer FB into both hindpaws labelled tibial afferents, before unilateral SNI_trans_ or SNI_crush_ surgery. (B) Representative image of FB labelling and NeuN immunostaining in the L4 DRG. Image is a projection of optical sections at 3 µm intervals through the entirety of a 30 µm-thick tissue section. Scale bar = 100 µm. (C-D) Quantification of cross-sectional area of FastBlue labelled DRG neurons ipsilateral and contralateral to SNI_trans_ (C) or SNI_crush_ injury (D) reveals a loss of small afferents and subsequent shift in population distribution. Kolmogorov-Smirnov tests of cumulative distributions; SNI_trans_: D = 0.25, *P* < 0.001; *n* = 183 or 191 neurons from 3 mice; SNI_crush_: D = 0.22, *P* < 0.001, *n* = 319 or 325 neurons from 3 mice. (E) Experimental approach for whole DRG volumetric analyses after SNI_trans_. (F) Representative 3D rendering of TDP-43 profiles and corresponding nuclear spot profiles following Imaris-based spot detection feature. Scale bar = 100 µm. (G) Quantification of DRG nuclear spot volume ipsilateral and contralateral to SNI_trans_. Kolmogorov-Smirnov tests of cumulative distribution: D = 0.06, *P* < 0.001, *n* = 30206 (contra) or 32544 (ipsi) nuclei from 4 (contra) or 5 (ipsi) mice. (H) Total number of nuclear spots, by size, per DRG. 2-way RM ANOVA; Size bin x Injury interaction: F_2,14_= 8.26, *P* = 0.004; *n* = 4-5 mice; Šídák’s multiple comparisons tests: ** *P* < 0.01.

As a third complementary approach, we applied semi-automated volumetric analyses of nuclei size following tissue clearing. Here, whole DRG were cleared 4 weeks after SNI_trans_ for nuclei counting in “complete” tissue (**Fig. 2E-H**). Nuclei were labelled by TDP-43, in line with [66] and were quantified using Imaris software (**Fig. 2F**, supplemental video 1). We observed a slight but significant rightward shift in nuclear spot volume population distribution 4 weeks after SNI_trans_ (**Fig. 2G**). Additionally, there was a significant reduction in the number of small but not medium or large nuclear spots, in support of a loss of small diameter neuron populations (**Fig. 2H**).

Together, our data derived from several different experimental approaches show that a population of small diameter afferents are lost following peripheral nerve injury.

### Spared nerve crush or transection results in death of Mrgprd-expressing neurons

To date, determining cell loss among specific populations of afferent neuron has proved challenging due to the downregulation of subpopulation-specific marker genes following axonal transection [36,43]. To overcome this issue, we took advantage of transgenic strategies to label populations in a manner that persisted after injury. Owing to the bias for loss of small neurons and the known loss of IB4-binding central terminals post-injury [35], we initially focussed on non-peptidergic nociceptive neurons. We used MrgD^ChR2-YFP^ mice to identify neurons belonging to the largest of the three classes of non-peptidergic nociceptors, NP1 [54,58]. To determine whether these neurons are lost following nerve injury, we used a stereological method to quantify L4 DRG MrgD-YFP^+^ neurons 28 days after sham surgery or SNI_trans_ (**Fig. 3A-B**). SNI_trans_, but not sham, resulted in a significant decrease (54.0 ± 6.6%) in the total number of MrgD-YFP^+^ neurons in L4 DRG (**Fig. 3C**).

**Figure 3.**
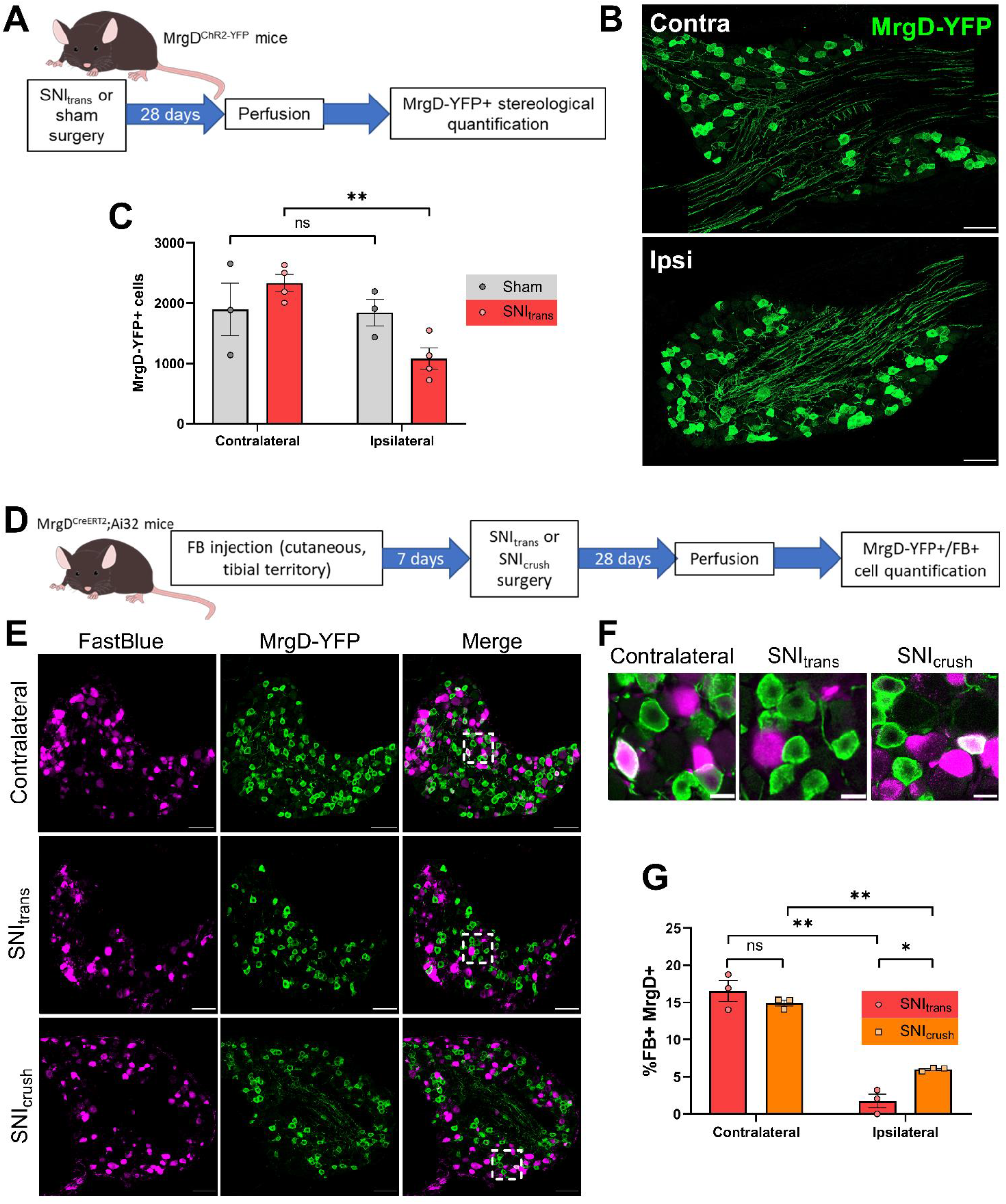
Spared nerve crush or transection results in death of non-peptidergic neurons. (A) Schematic of experimental approach for (B) and (C). (B) MrgD^ChR2-YFP^ L4 DRGs 4 weeks after SNI, contralateral or ipsilateral to injury. Images are projections of optical sections at 3 µm intervals through the entirety of 30 µm-thick tissue sections. Scale bars = 100 µm. (C) Quantification of total number of MrgD-YFP+ cells per L4 DRG 4 weeks after SNI revealed a significant loss in ipsilateral DRG. 2-way RM ANOVA with Šídák’s multiple comparisons tests; Side x Treatment interaction: F_1,5_ = 9.23, *P* = 0.029; *n* = 3 mice. (D) The experimental approach used to generate data presented in (E-G). (E-F) MrgD-YFP expression and FB labelling in the L4 DRG, 14 days after SNI or crush surgery, or contralateral to injury. White boxes represent regions enlarged in (F). Scale bars = 100 µm (E) or 20 µm (F). (G) The proportion of FB-labelled DRG neurons decreased after spared nerve crush injury, and co-labelling is almost completely absent after SNI. 2-way RM ANOVA with Šídák’s multiple comparisons tests; Side x Injury interaction: F_1,4_ = 7.80, *P* = 0.049; *n* = 3 mice. Post-tests: * *P* < 0.05, ** *P* < 0.01.

YFP expression in MrgD^ChR2-YFP^ mice is driven by the endogenous *Mrgprd* promotor, which has been reported to be either up-or downregulated following axonal damage [43,57]. Such changes in promoter activity could affect the proportion of non-peptidergic nociceptors identified by YFP expression. Therefore, to verify these findings, we used MrgD^CreERT2^;Ai32 mice and tamoxifen administration prior to injury, to permanently label *Mrgprd-*expressing afferents with ChR2-YFP (**Fig. 3D-F**). We then tested whether the proportion of cutaneous tibial afferents that were YFP^+^ was altered following nerve injury. Following hindpaw FB injection, ∼15% of contralateral, FB-labelled DRG neurons expressed YFP. This was reduced to 6.0 ± 1.2% 28 days after SNI_crush_ injury, and to only 1.7 ± 0.9% 28 days after SNI_trans_ (**Fig. 3G**). Uptake by uninjured YFP^+^ neurons was equivalent 7 and 35 days after FB injection, demonstrating that this reduction was not because 7 days was insufficient for YFP^+^ neurons to fully uptake FB (**Fig. S2C**). No significant difference in the percentage of FB-labelled YFP^+^ DRG neurons between ipsilateral and contralateral DRG was observed at 7 days following SNI_trans_ (**Fig. S3A-B**), demonstrating that loss occurred after this timepoint. Collectively, these data show that peripheral nerve injury results in a substantial loss of non-peptidergic, *Mrgprd*-expressing neurons, with SNI_trans_ (i.e., an unrepaired axonal transection) resulting in an almost complete loss of this population.

### Spared nerve injury induces a loss of Trpm8^+^ and CGRP^+^ but not myelinated DRG neurons

Loss restricted to non-peptidergic nociceptors would not fully account for the degree of total neuron loss that we observed. Therefore, we studied a range of other sub-populations, both small and large in diameter, for their vulnerability to injury-induced loss. To investigate potential loss of Trpm8^+^ (cold-sensitive), CGRP^+^ (peptidergic), and myelinated subpopulations of DRG neurons following nerve injury, we applied our FB-labelling approach in Trpm8^FlpO^;RC::FLTG (FlpO-dependent tdTom expression), Calca^CreERT2^;Ai32 (Cre-dependent ChR2-YFP expression) and Thy1-CFP mice respectively (**Fig. 4A-D**). Trpm8-tdTom was expressed by a population of small-diameter, putative cold-sensitive neurons (**Fig. 4B**), accounting for 8.3 ± 0.27% of FB-labelled neurons in contralateral DRG. This decreased to 4.2 ± 0.96% ipsilateral to SNI_trans_ injury (Fig. 4E), indicating a partial loss of Trpm8^+^ afferents. When examining peptidergic afferents, we found that 48.1 ± 2.42% of FB-labelled neurons in contralateral DRG were Calca-YFP^+^, compared to 34.3 ± 2.54% 4 weeks after SNI_trans_ injury (Fig. 4C, F), consistent with a partial loss of CGRP^+^ afferents. We used a Thy1-CFP line that demonstrates consistent expression post-injury [60] and labels a sample of medium/large diameter myelinated afferents. CFP was largely restricted to NF200^+^ neurons, labelling 56% of this population. Expression was present in a heterogenous population of nociceptive (TrkA+) and non-nociceptive (TrkA-) myelinated neurons (**Fig. S4**). Contralateral to injury, 15.6 ± 1.8% of FB-labelled neurons expressed Thy1-CFP (Fig. 4D, G). In contrast to unmyelinated subpopulations, this proportion was higher in ipsilateral DRG following SNI_trans_ (23.3 ± 3.2%), consistent with no (or minimal) loss of Thy1-CFP-expressing afferents, accompanied by a loss of Thy1-CFP-negative neurons. Collectively, these data show that unrepaired axonal damage to peripheral sensory neurons induces a partial loss of Trpm8^+^ and CGRP^+^ subpopulations, but no major loss of myelinated afferents.

**Figure 4.**
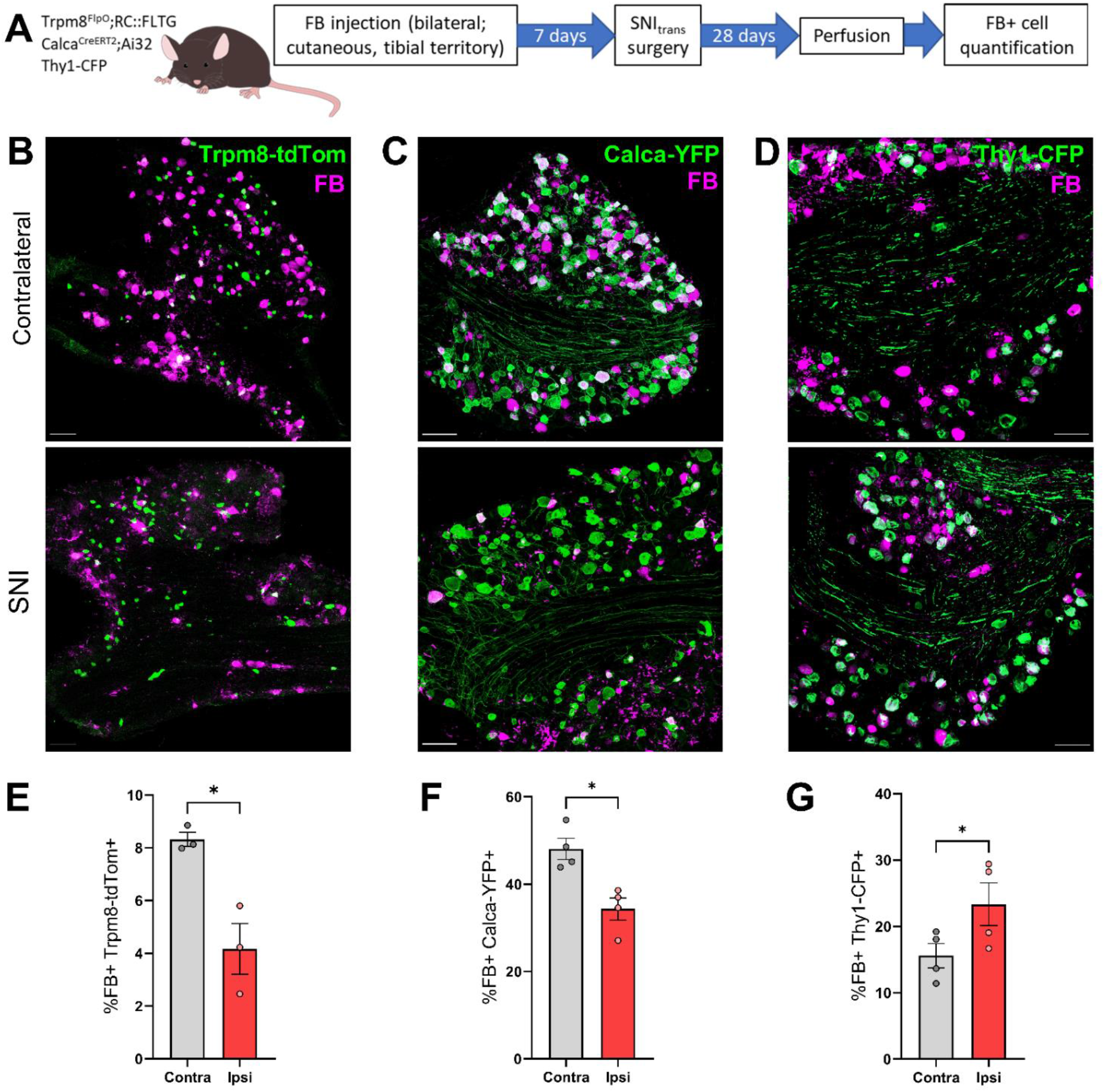
Spared nerve injury induces a loss of Trpm8+ and CGRP+ but not myelinated DRG neurons. (A) Schematic of experimental approach. (B-D) FB labelling and Trpm8-tdTom (B), Calca-YFP (C) or Thy1-CFP expression (D) 28 days after SNI_trans_ in the L4 DRG, contralateral (top) or ipsilateral (bottom) to injury. Images are projections of optical sections at 3 µm intervals through the entirety of 30 µm-thick tissue sections. Scale bars = 100 µm. (E-G) Quantification of the proportion of FB-labelled neurons also expressing Trpm8-tdTom (E), Calca-YFP (F) or Thy1-CFP (G) in L4 DRG contralateral or ipsilateral to SNI_trans_. Paired t-tests; Trpm8-tdTom: t_2_ = 5.31, *P* = 0.034, *n* = 3 mice; Calca-YFP: t_3_ = 4.12, *P* = 0.026, *n* = 4 mice; Thy1-CFP: t_3_ = 4.42, *P* = 0.022, *n* = 4 mice. * *P* < 0.05.

Based upon our finding of preferential loss of non-peptidergic nociceptors, we re-analysed a previous population-specific transcriptomic dataset of mouse DRG neurons following nerve injury for potential upregulation of cell death pathways (**Fig. S5**) [3]. We found that early after injury (3 days post-SNI_trans_), non-peptidergic (MrgD^CreERT2^-expressing) neurons showed enhanced enrichment of GO terms associated with apoptosis, in contrast to a broad population of nociceptors (labelled with Scn10a^CreERT2^), peptidergic nociceptors (Calca^CreERT2^), C-LTMRs (Th^CreERT2^), and Aβ-RA (rapidly adapting) and Aδ-LTMRs (Aδ/Aβ-LTMR, Ntrk2^CreERT2^;Advillin^FlpO^), in which there was less or no enrichment of cell death pathways. By 4 weeks, only C-LTMR and Aδ/Aβ-LTMR subtypes show any over-representation of cell death pathways (in the populations studied). Both injury-specific and apoptotic signatures in non-peptidergic neurons were no longer significantly enriched, consistent with a loss of axotomized non-peptidergic afferents by this late timepoint post-injury. These data suggest that apoptotic pathways are upregulated acutely after injury in a cell-type specific manner.

### Mrgprd DRG neurons are sensitive to loss *in vitro*

Earlier studies postulated that a lack of neurotrophic support underlies neuronal loss, which is supported by the observation that exogenous GDNF treatment at the time of injury, or shortly after, rescues the loss of IB4-binding central terminals post-transection [5]. We sought to use the DRG neurons from MrgD^CreERT2^;Ai32 mice to test this postulate and establish an *in vitro* platform capable of probing the molecular basis of loss, with axonal transection during isolation providing a correlate for *in vivo* nerve injury (**Fig. 5A-E**). 24-hours after plating, YFP was expressed by 16.3 ± 1.3% of DRG neurons, which was reduced to 11.8 ± 1.7% after 28 days of culture in the presence of exogenous growth factors (GFs), NGF and GDNF (**Fig. 5F**). However, in the absence of GFs, YFP^+^ neurons only accounted for 1.7 ± 0.6% of neurons after 28 days, accompanied by an apparent reduction in the overall number of neurons within the culture, despite all conditions being seeded at the same initial density (**Fig. 5C, F**). YFP^+^ cell loss was partially rescued by the presence of GDNF, but not NGF alone, in the culture media (**Fig. 5 D-F**). Collectively, these data support the use of DRG cultures to probe the mechanisms underlying loss of sensory neurons following nerve injury, and suggest a role for trophic support, particularly via GDNF signalling, in preventing the loss of non-peptidergic nociceptors.

**Figure 5.**
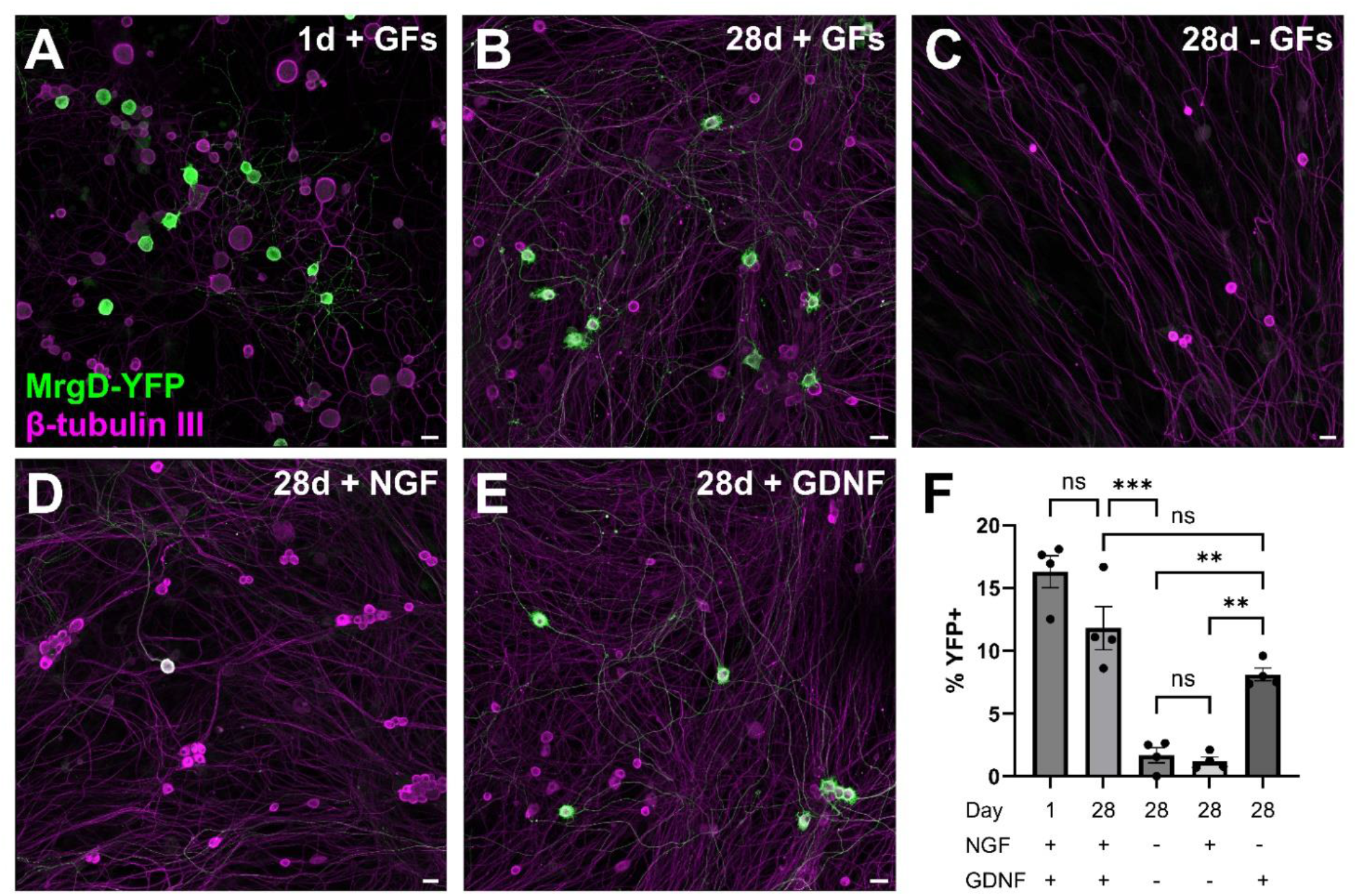
Neurotrophic support ameliorates MrgD+ cell loss *in vitro.* (A-E) Representative fields of view of MrgD-YFP and β-tubulin III expression in neuronal cultures of isolated MrgD^CreERT2^;Ai32 DRGs, 24 hours (A) or 4 weeks (B-E) after plating, with the addition of the growth factors (GFs) NGF, GDNF, both, or neither. Scale bars = 20 µm. (F) Quantification of the percentage of neurons that express MrgD-YFP under the conditions shown in (A). One-way RM ANOVA with Tukey’s post-tests; F_4_,_10_ = 51.6, *P* < 0.001; *n* = 4 mice. Post-tests: * *P* < 0.05, ** *P* < 0.01, *** *P* < 0.001.

## Discussion

We present data herein to support the hypothesis that traumatic nerve injury in rodents leads to a profound loss of small-diameter DRG neurons. Taking advantage of newly developed transgenic recombinase driver lines, we have shown that loss is biased across molecularly defined sub-populations. Non-peptidergic nociceptive neurons are particularly susceptible to loss, with almost all Mrgprd^+^ axotomized afferents lost following an unrepaired transection injury (SNI_trans_) and roughly half lost following a model which contrastingly allows for nerve regenerations (SNI_crush_). Finally, we have observed that the vulnerability of Mrgprd^+^ neurons extends to the *in vitro* setting and provide data to support the hypothesis that loss is driven by a lack of neurotrophic support following injury.

### Neuronal loss

The question of whether DRG neurons die following traumatic injury has been addressed by several groups over the last few decades. Despite contrasting findings on the extent, timing, and form that loss takes, most studies have observed frank loss of DRG neurons [6,37,45,52]. However, more recent studies using recombinase driver lines and novel machine-learning approaches have cast doubt on this consensus [43,48]. Our data strongly supports the loss hypothesis and suggests that around 60% of axotomized afferents die within two weeks of SNI. The discrepancy between our findings and other recent studies may be partly explained by the sampling method used to estimate neuronal number. For example, Shulte *et al* developed a novel machine-learning approach and found no reduction in neuron density across serial sections of rat DRG following SNI, and inferred from this that frank loss did not occur [48]. Our results are congruous, in that we also observed no reduction in neuron density. However, we found a substantial loss in the total neuron-containing volume of injured DRG, which underlies our contrasting conclusion of frank loss. Of note, morphological volumetric analysis and MRI have also previously demonstrated volume loss in both rodent and human DRG following nerve injury [34,64,65]. These findings occur despite a major increase of non-neuronal cells in the injured DRG [30] and support the notion that total DRG neuron number is decreased.

### Selectivity of neuron loss

While definitively characterising loss of molecularly defined subpopulations was challenging before the advent of recombinase driver lines, a consensus emerged that small diameter neurons are more vulnerable to nerve injury-induced loss [49,52]. Our data support this consensus and extend it to reveal that while there is a generalised partial loss of C-fibre populations including CGRP-and Trpm8-expressing neurons, Mrgprd-expressing neurons are particularly sensitive to loss. This selective vulnerability has been hinted at previously by the stark reduction in the number of DRG neurons and their central terminals that bind IB4 and express canonical markers such as P2X_3_ receptor following nerve injury [5,8,29,35]. Type 1a glomeruli are also reduced in lamina II, suggesting a structural loss of central terminals and not simply a loss of IB4-binding [2]. However, it was not clear whether these data represented phenotypic changes in non-peptidergic nociceptors or frank loss of neurons. We describe neuron loss that is delayed (occurring >7 days post-injury) in respect to histochemical and structural changes (occurring 1-5 days post-injury [2,29]), suggesting that these changes precede and are not in themselves indicative of neuron loss.

The vulnerability of Mrgprd-expressing neurons is congruous with recent subpopulation bulk RNAseq data, which found that SNI-related gene expression signatures were less evident in Mrgprd-expressing and C-LTMR neurons at later timepoints, compared to other populations in injured DRG [3]. This could be explained by a loss of axotomized neurons of these classes and therefore sampling of only uninjured neurons at this timepoint. Such vulnerabilities could also explain previously identified differences based on functional type. Nerve transection results in loss of cutaneous but not muscle afferents [24,63]. Non-peptidergic nociceptive neurons make up a far greater proportion of cutaneous afferents compared to muscle afferents (32.9 vs 3.7% when comparing sural and medial gastrocnemius afferents in rat [42]), which could explain the differing vulnerabilities. In terms of transcriptional response to injury, non-peptidergic nociceptors show enrichment of individual pro-apoptotic factors early after injury [23,67], and we extend these results here, by describing a subpopulation specific enrichment of GO terms associated with apoptosis that is evident as early as three days after injury. Such data, and single cell transcriptomic profiling of all DRG neurons following injury [36,43], may offer the opportunity to elucidate the cell death pathways engaged and upstream effectors that enrich this process to non-peptidergic nociceptive neurons.

### Implications for pain pathogenesis

Neuronal loss has been proposed as a key contributor to poor functional recovery following nerve injury [53] and biased survival of different afferent types might be expected to contribute to modality-specific sensory deficits. Beyond loss of function, does DRG neuron loss contribute to chronic pain, in either an adaptive or maladaptive manner? Intrathecal delivery of GDNF is neuroprotective and reverses the reduction in the number of IB4-binding DRG neurons and central terminals seen following transection [5]. Treatment is concurrently analgesic and abrogates pain-related behaviours [7,59]. However, the pleiotropic nature of GDNF makes it impossible to directly attribute the analgesic effects to reversal of neuron loss. Indeed, it is possible that GDNF exerts its effect via actions on intact non-peptidergic nociceptive afferents [51], activation of which is known to drive aversive behaviours in the neuropathic state [61]. These data leave the contribution of non-peptidergic nociceptor loss to behaviour in the GDNF treatment paradigm ambiguous. Other pharmacological approaches have been found effective at reversing neuronal loss in rodent models, but the impact on pain behaviour was not studied [21,22].

Rodents develop marked mechanical and thermal hypersensitivity rapidly following nerve injury, and before timepoints at which neuron loss is observed [10]. This lack of a temporal correlation may suggest a limited contribution to evoked hypersensitivities. The temporal profile of ongoing tonic pain (for example, pain aversiveness as measured by condition place preference assays [26]) is less defined, and therefore so is its correlation to the timing of neuron loss.

There are many anatomical sites within the somatosensory nervous system where differential loss of sensory neuron populations could impact neurobiology. For example, loss of cutaneous afferents may afford more opportunity for plasticity in reinnervation patterns, such as collateral sprouting of uninjured or surviving afferents, and the types of nerve endings made by different molecular subpopulations [17,27]. It also seems likely that death of many neurons within a DRG could contribute to the expansion and activation of immune cell types, which is known to play a major role in neuropathic pain [30,69]. Finally, under normal conditions, peripheral sensory input is integrated in the dorsal horn of the spinal cord by complex interneuron circuitry. Many spinal circuits are engaged by convergent input from different afferent types [9,40,70]. Therefore, selective loss of input from discrete afferent types could undoubtedly impact the normal processing of remaining afferent signals [33]. Experimentally abrogating neuronal loss may be a fruitful approach to assess the contribution to nervous system plasticity (adaptive or maladaptive) following injury. In this regard, our *in vitro* readout would be a useful experimental platform to help delineate the precise cell death pathways and signalling cascades engaged (and which could then be experimentally manipulated). Such studies should consider that plasticity may evolve over time. The loss of IB4^+^ central terminals is transient following crush and has even been observed to reverse at longer timepoints following SNI_trans_ [35]. These observations, in conjunction with ours of loss of neurons, raises the intriguing question of the source of such central reinnervation.

### Study limitations

Our efforts focussed on traumatic nerve injury paradigms owing to previous contrasting results using these robust and reproducible experimental models. We did not extend our studies to systemic neuropathy models, such as those modelling chemotherapy or diabetic neuropathy. A recent post-mortem analysis reported neuronal loss in the DRG from patients who had been experiencing painful diabetic peripheral neuropathy [19]. Transcriptional responses vary substantially across different nerve insults [43] and so it would be of interest to test whether neuronal loss and the subpopulation vulnerability reported here are common features across different types of insults.

Whilst we observed a frank loss of small-diameter neurons using multiple approaches, the extent of loss observed using semi-automated quantification in cleared, whole DRG was not to the same extent as that observed using manual techniques, nor were we able to replicate the general cell loss after SNI_trans_ documented by West et al (2020) using cleared tissue [66]. Two major limitations here may explain this discrepancy: First, due to technical issues, the dataset is unpaired ipsilateral-contralateral which add larger variability. Second, the analysis method is prone to undercounting deep nuclei. The signal-to-noise is better for superficial nuclei and smaller tissue volumes. Given the reduction in DRG volume after SNI_trans_, nuclei in larger contralateral DRG may be undercounted.

While we made efforts to profile loss of several molecularly discrete sensory neuron populations, we acknowledge that not all subtypes were profiled. Furthermore, recent single cell RNA sequencing has given us a more granular appreciation of the heterogeneity of sensory neurons, and excitingly, new recombinase lines have been developed to genetically target these subpopulations [41]. Future studies could leverage our experimental approach and these new transgenic lines to characterise the loss of neurons in more detail. Such experiments may be pertinent before embarking on molecular or functional profiling of populations post-nerve injury.

### Conclusions

In sum, we have provided data from multiple complimentary experimental approaches to support the hypothesis that DRG neurons are lost following nerve injury in mice. We describe a substantial loss, which is biased towards specific subpopulations and particularly present in small diameter non-peptidergic nociceptive neurons.

## Acknowledgments

We thank Dr Mark Hoon for providing the Trpm8-Flp transgenic mouse line, and Prof Andrew Todd and Dr David Hughes for their critical feedback on the manuscript. Research was funded by an MRC Fellowship grant awarded to G.A.W. (MR/T01072X/1) and a Tenovus Scotland Pilot Grant awarded to A.H.C. and G.A.W. (S22-17). This work was also funded by the Wellcome Trust (DPhil scholarship to AMB, 215145/Z/18/Z) and a Wellcome Investigator Grant to DB (223149/Z/21/Z), as well as the MRC (MR/T020113/1), and with funding from the MRC and Versus Arthritis to the PAINSTORM consortium as part of the Advanced Pain Discovery Platform (MR/W002388/1). AMB further received a GTC MSDTC Scholarship.

DB has acted as a consultant in the last 2 years for AditumBio, Biogen, Biointervene, Combigene, LatigoBio, GSK, Ionis, Lexicon therapeutics, Neuvati, Olipass, Orion, Replay, SC Health Managers, Theranexus, Third Rock ventures, Vida Ventures on behalf of Oxford University Innovation. DB has received research funding from Lilly and Astra Zeneca and GW has received research funding from Ono Pharmaceutical. DB has received an industrial partnership grant from the BBSRC and AstraZeneca.

Other authors have no conflicts of interest to declare.

---

**Figure S1 (Data related to Figure 1).**
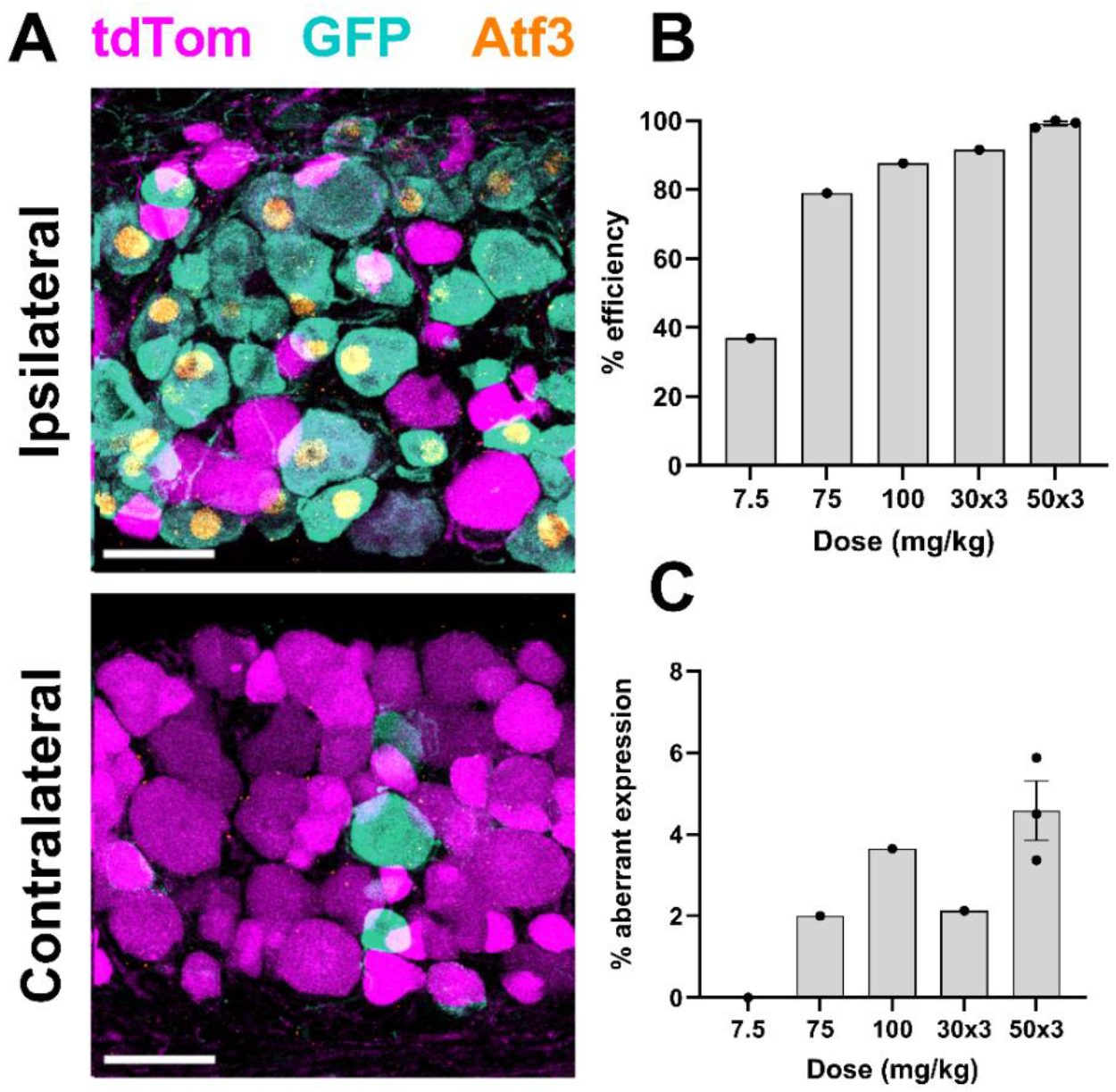
Tamoxifen dose-dependently increases recombination efficiency in Avil^FlpO^;Atf3^CreERT2^;RC::FLTG mice. (A) tdTomato (uninjured), GFP (injured) and Atf3 expression 14 days after SNI_trans_ surgery at the ipsilateral and contralateral L4 DRG, following 50 mg/kg tamoxifen at 0, 3 and 7 days post-SNI, the dosing regimen used when collecting data presented in Figure 1. Images are projections of optical sections at 3 µm intervals through the entirety of 30 µm-thick tissue sections. Scale bars = 50 µm. (B) Percentage efficiency of Atf3 recombination at the ipsilateral DRG (i.e., the percentage of Atf3-expressing neurons that co-express GFP) at the initial pilot dosing regimens tested for efficiency of recombination. *n* = 1-3 mice. (C) Quantification of aberrant expression of GFP at the contralateral DRG (i.e., the percentage of neurons expressing GFP).

**Figure S2 (related to Figures 2 to 4).**
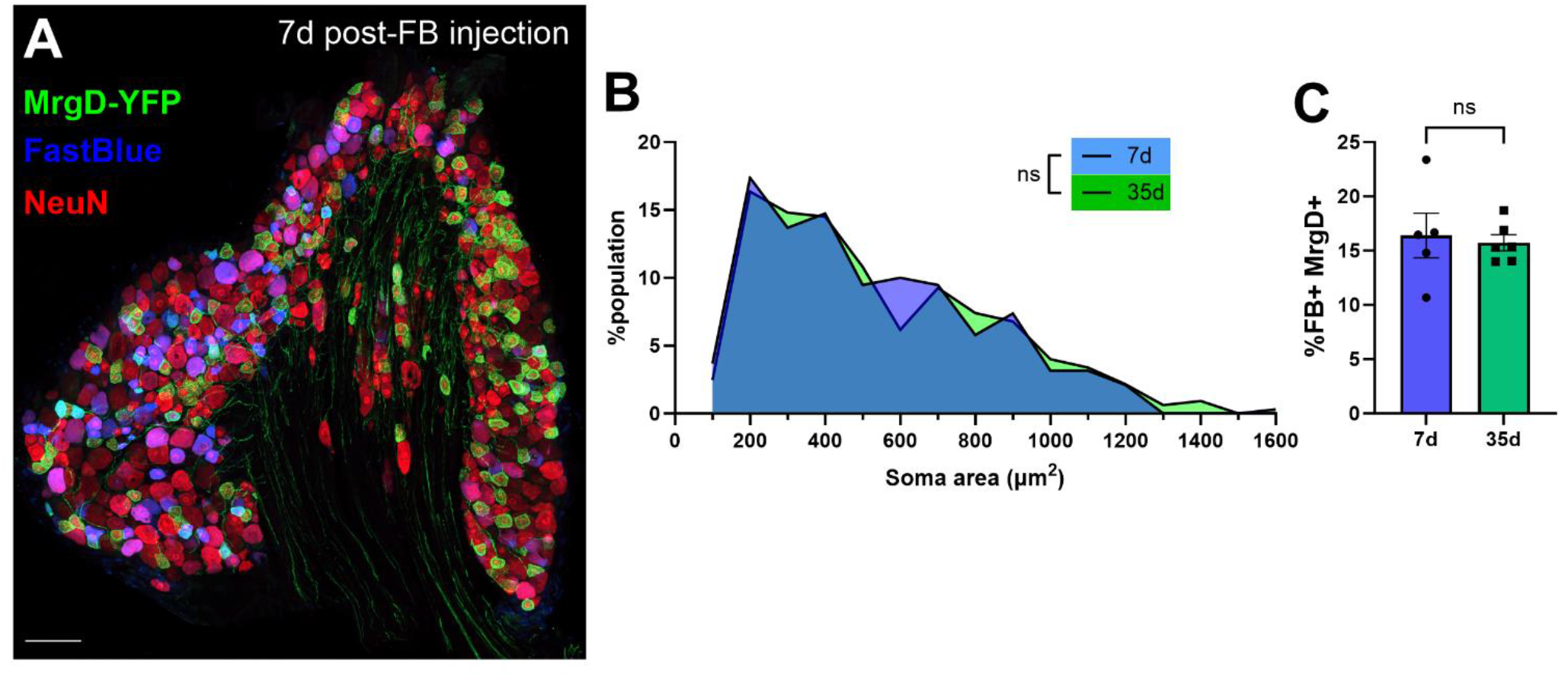
**FastBlue (FB) uptake is equal at 7- and 35-days post-injection**, i.e., the potential for reduced FB uptake at 7D vs 35D after injection is not responsible for differing populations of FB-filled cells observed in the ipsilateral versus contralateral DRG following SNI_trans_. (A) FB labelling and MrgD-YFP and NeuN expression in the L4 DRG of a MrgD^CreERT2^;Ai32 mouse 7 days post-FB injection. Image is a projection of optical sections in the Z axis at 3 µm intervals through the entirety of a 30 µm tissue section. (B) Population distributions of FB-labelled neurons 7- or 35-days after FB injection. Kolmogorov-Smirnov test: D = 0.057, *P* = 0.83, *n* = 191-324 neurons from 5-6 mice. (C) Quantification of the percentage of FB-labelled neurons in the L4 DRG that are MrgD-YFP+ 7- or 35-days post-FB injection. 35-day data is pooled from the DRGs contralateral to SNI_trans_ or SNI_crush_ presented in Figure 3G. Unpaired t-test; t_9_ = 0.34, *P* = 0.75, *n* = 5-6 mice.

**Figure S3 (related to Figure 3).**
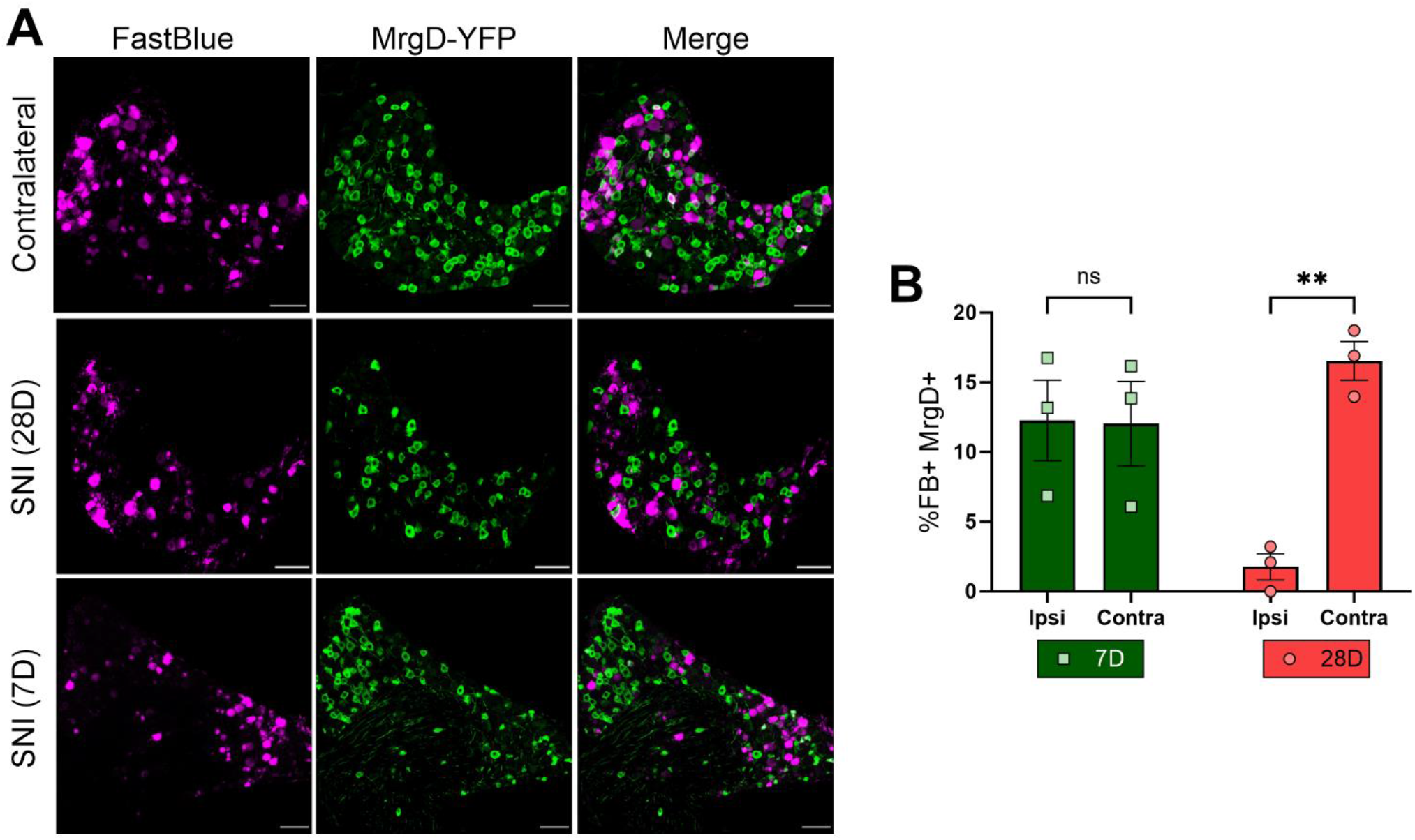
Non-peptidergic neuron death does not occur until after 7d post-nerve injury. (A) FB labelling and MrgD-YFP expression in MrgD^CreERT2^;Ai32 L4 DRGs 7- or 28-days after SNI_trans_ surgery, or contralateral to injury. 28-day post-SNI_trans_ data also shown in Fig. 3. Scale bars = 100 µm. (B) Quantification of the percentage of FB-labelled neurons that are MrgD-YFP+. 2-way RM ANOVA; Timepoint x Side interaction: F_1,4_ = 51.3, *P* = 0.002; Šídák’s multiple comparisons tests: ** *P* < 0.01; *n* = 3 mice.

**Fig S4 (related to Figure 4).**
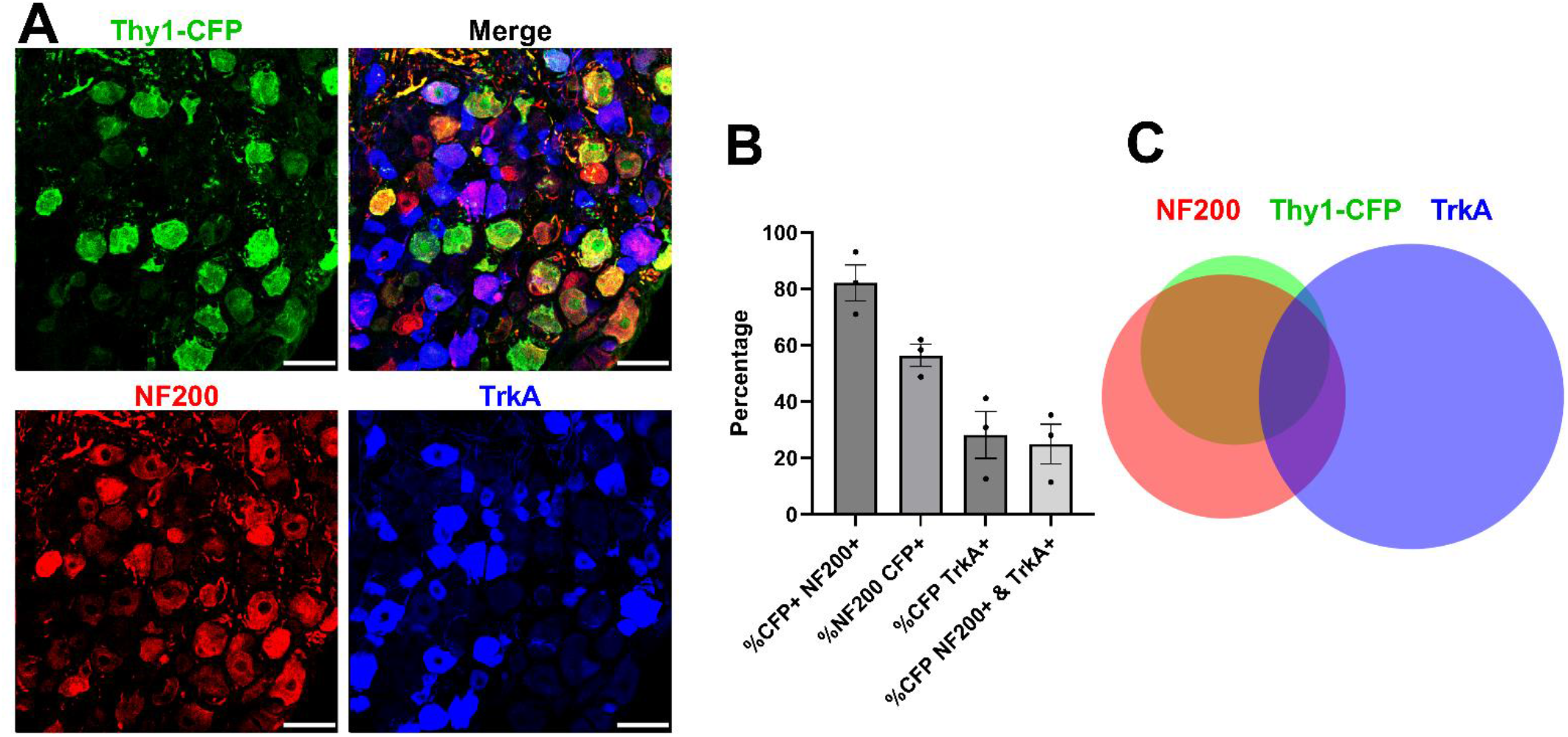
Thy1-CFP is expressed by a heterogenous population of myelinated afferents. (A) expression of Thy1-CFP, NF200 (expressed by myelinated afferents) and TrkA (expressed by a majority of peptidergic afferents) in a lumbar DRG. Image is a projection of optical sections in the Z axis at 3 µm intervals through the entirety of a 30 µm tissue section. (B) Quantification of overlap of expression. *n* = 3 mice. (C) Quantitative Venn diagram illustrating overlap of expression; *n* = 1503 neurons from 3 mice.

**Figure S5 (related to Figure 4).**
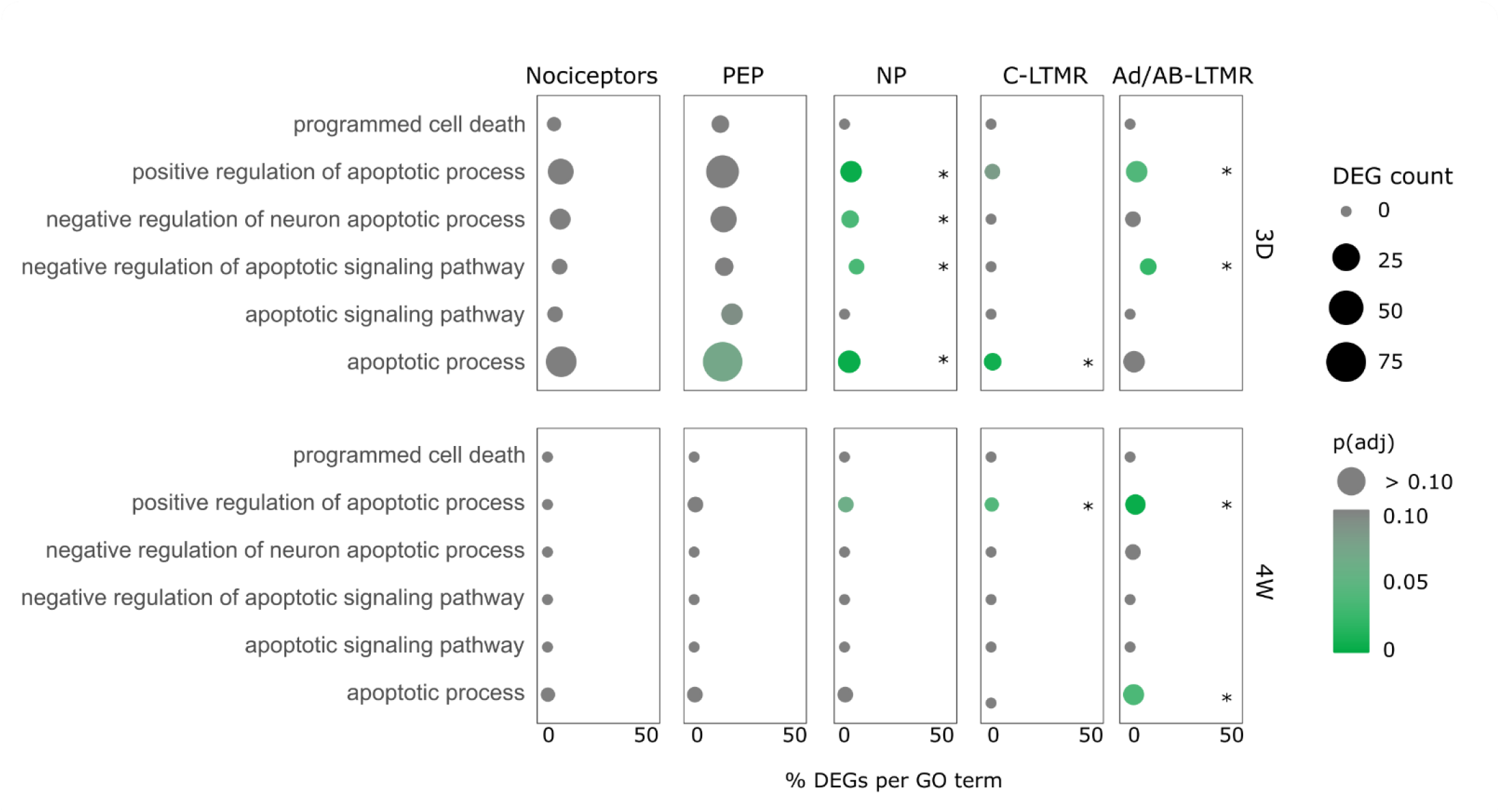
Non-peptidergic neurons show and overrepresentation of cell death pathways 3 days after SNI_trans_. Gene Ontology (GO) term analysis for cell death pathways across DRG subtypes after SNI (Barry et al., 2023). Five subpopulations were included at 3 days (3D) and 4 weeks (4W) after surgery: general nociceptors (labelled with Scn10a^CreERT^), peptidergic nociceptors (PEP, Calca^CreERT2^), non-peptidergic nociceptors (NP, Mrgprd^CreERT2^), C-LTMRs (Th^CreERT2^), and Aß-RA (rapidly adapting) + Aδ-LTMRs (Aδ/Aβ-LTMR, Ntrk2^CreERT2^;Avil^FlpO^). Significantly over-represented terms (*P* < 0.05) per population and timepoint are represented with an *. DEG, differentially expressed genes.

